# PIP4Ks Suppress Insulin Signaling Through a Catalytic-Independent Mechanism

**DOI:** 10.1101/370544

**Authors:** Diana G. Wang, Marcia N. Paddock, Mark R. Lundquist, Janet Y. Sun, Oksana Mashadova, Solomon Amadiume, Timothy W. Bumpus, Cindy Hodakoski, Benjamin D. Hopkins, Matthew Fine, Amanda Hill, T. Jonathan Yang, Jeremy M. Baskin, Lukas E. Dow, Lewis C. Cantley

## Abstract

Insulin stimulates conversion of phosphatidylinositol-4,5-bisphosphate (PI(4,5)P_2_) to phosphatidylinositol-3,4,5-trisphosphate (PI(3,4,5)P_3_), which mediates downstream cellular responses. PI(4,5)P_2_ is produced by phosphatidylinositol-4-phosphate 5-kinases (PIP5Ks) and byphosphatidylinositol-5-phosphate 4-kinases (PIP4Ks). Here we show that deletion of the three genes that encode PIP4Ks (*PIP4K2A, PIP4K2B* and *PIP4K2C*) *in vitro* results in a paradoxical increase in PI(4,5)P_2_ and a subsequent increase in insulin-stimulated production of PI(3,4,5)P_3_. Surprisingly, reintroduction of either wild-type or kinase-dead forms of the PIP4Ks restored cellular PI(4,5)P_2_ levels and insulin stimulation of the PI3K pathway. These effects are explained by an increase in PIP5K activity upon deletion of PIP4Ks, which we demonstrate can suppress PIP5K activity *in vitro* through a direct binding interaction. Collectively, our work reveals an important non-catalytic function of PIP4Ks in suppressing PIP5K-mediated PI(4,5)P_2_ synthesis and insulin-dependent conversion to PI(3,4,5)P_3_ by PI3K enzymes and suggests that pharmacological depletion of PIP4K enzymes using emerging degrader technologies could represent a novel strategy for stimulating insulin signaling.

## INTRODUCTION

Phosphatidylinositol-4,5-bisphosphate (PI(4,5)P_2_) plays numerous roles in cellular regulation. It mediates actin remodeling at the plasma membrane and plays multiple roles in vesicle trafficking. It is the substrate that hormone-stimulated phospholipases type C utilize to generate the second messengers diacylglycerol and inositol-1,4,5-trisphosphate (Balla, 2013; Sun et al., 2013). It is also the substrate that Class 1 phosphoinositide 3-kinases utilize to generate the second messenger phosphatidylinositol-3,4,5-trisphosphate (PI(3,4,5)P_3_) in response to insulin and other growth factors (Fruman et al., 2017).

Mammals have three genes that encode enzymes that generate PI(4,5)P_2_ from phosphatidylinositol-4-phosphate (PI(4)P) and three genes that generate PI(4,5)P_2_ from phosphatidylinositol-5-phosphate (PI(5)P) (Rameh et al., 1997). The former three genes are named *PIP5K1A*, *PIP5K1B* and *PIP5K1C*, and we will refer to the class of enzymes they encode as PIP5Ks (also known as PI-4-P5Ks, Type 1 PIP kinases, or PIP5Kα/β/γ) (van den Bout and Divecha, 2009). The latter three genes are named *PIP4K2A*, *PIP4K2B* and *PIP4K2C*, and they encode PIP4Ks (also known as PI-5-P4Ks, Type 2 PIP kinases, or PIP4Kα/β/γ). Yeast have only one enzyme for generating PI(4,5)P_2_, encoded by *MSS4*, a phosphatidylinositol-4-phosphate 5-kinase (PIP5K) (Homma et al., 1998). However, multicellular animals from flies and worms to mammals have genes from both families. The enzymes encoded by these two families of genes have sequence and structural similarities, and a single amino acid change in the phosphoinositide substrate binding pocket can change a PIP4K into a PIP5K (Kunz et al., 2001; Kunz et al., 2000).

PI(4)P is over 100-fold more abundant than PI(5)P in cells, and it is generally assumed that most PI(4,5)P_2_ in mammalian cells is generated from PI(4)P via PIP5Ks (Balla, 2013). In cell lines and tissues where phosphoinositides have been quantified, PI(4)P and PI(4,5)P_2_ are at similar levels. Whereas local levels of PI(4,5)P_2_ can transiently drop when cells are stimulated with growth factors or hormones that activate phospholipases C or phosphoinositide 3-kinases, the total levels of PI(4,5)P_2_ and PI(4)P remain remarkably constant, indicating tight homeostatic control of these lipids.

Because PI(5)P is far less abundant than PI(4)P and is thought to contribute little to total cellular PI(4,5)P_2_ synthesis, there has been speculation that the function of the PIP4Ks is to decrease the level of PI(5)P (Jones et al., 2006; Wilcox and Hinchliffe, 2008). The bulk of PI(5)P appears to be generated in a two-step process. First, the lipid kinase PIKFYVE converts PI(3)P to PI(3,5)P_2_ for the purpose of maturing early endosomes into late endosomes and lysosomes. Second, a myotubularin family 3-phosphatase converts PI(3,5)P_2_ to PI(5)P (Jefferies et al., 2008; Zolov et al., 2012). This latter step probably occurs at the late endosome/lysosome, though this process is difficult to visualize due to the low abundance of PI(5)P and lack of selective probes for these lipids (Idevall-Hagren and De Camilli, 2015). A recent publication from our laboratory showed that conversion of PI(5)P to PI(4,5)P_2_ by PIP4Kα/β, likely on lysosomes, is a critical step in mediating fusion between autophagosomes and lysosomes (Lundquist et al., 2018). This study argued that while the PIP4Ks generate only a small fraction of cellular PI(4,5)P_2_, the location of the PI(4,5)P_2_ generated by these enzymes plays a critical role in completion of autophagy in some tissues. There is also evidence for pools of PIP4Kα/β and its substrate PI(5)P enzymes in the nucleus, so the PIP4K enzymes are likely to have many other functions beyond completion of autophagy (Bultsma et al., 2010; Jones et al., 2006).

Homozygous germline deletion of any one of the three murine PIP4Ks results in a viable mouse with normal development into adulthood (Emerling et al., 2013; Lamia et al., 2004; Shim et al., 2016). Interestingly, *Pip4k2b^-/-^* mice exhibit reduced adiposity and increased insulin sensitivity in muscle (Lamia et al., 2004), and *Pip4k2c^-/-^* mice exhibit enhanced TORC1 signaling and autoimmune disease that could be reversed by the TORC1 inhibitor rapamycin (Shim et al., 2016). Importantly, these two studies indicated that PIP4K family members could suppress insulin/TORC1 signaling *in vivo*; however, the mechanism by which this occurred was not clear.

In this study, we sought to determine the mechanism by which PIP4Ks are regulating insulin and TORC1 signaling. To do this we removed PIP4Ks from multiple cell lines using either CRISPR or tandem miRNA-based short hairpin RNAs. We find that in the absence of all PIP4K enzymes there is a paradoxical increase in PI(4,5)P_2_ and decrease in PI(4)P due to an increase in PIP5K activity. Importantly, we show that control of PIP5K activity by PIP4Ks is not dependent on their catalytic function expression of endogenous levels of either kinase-active or kinase-dead (PIP4K^KD^) forms of PIP4Ks restores normal phosphoinositides levels and insulin signaling. Finally, we also show that the increase in PI(4,5)P_2_ is concomitant with enhanced insulin-dependent PI(3,4,5)P_3_ production and downstream AKT activation. Thus, we demonstrate that PIP4Ks have distinct catalytic and non-catalytic functions in controlling cellular metabolism. Furthermore, it is the loss of catalytic-independent functions of PIP4Ks that underlie the enhanced insulin and TORC1 signaling in *Pip4k2b^-/-^* and *Pip4k2c^-/-^* mice.

## RESULTS

### Loss of PIP4K family members does not affect cell viability

To investigate the role of PIP4K enzymes in cellular signaling, we generated tools to systematically deplete individual members of the PIP4K family. First, we optimized a series of tandem miRE-based short hairpin RNAs (shRNAs) to knockdown the PIP4Ks (Table S1, Table S2). We then expressed these from a constitutive lentiviral construct to generate cell lines with stable knockdown of PIP4K family isoforms in HeLa cells, as well as other human immortalized or transformed cells (Figure 1A, Figures S1A-C, Figures S2A-E) (Fellmann et al., 2013; Fellmann et al., 2011; Pelossof et al., 2017). shRNA-mediated silencing of one or all of the PIP4Ks did not result in gross changes in morphology or cause a major change in growth rate of the cells examined (Figure S1D). Second, as an orthogonal approach, we used CRISPR/Cas9-mediated mutagenesis to delete *PIP4K2A, PIP4K2B*, and *PIP4K2C* in HEK293T cells (Figure 1B, Table S3), with similar results (Ran et al., 2013).

**Figure 1.**
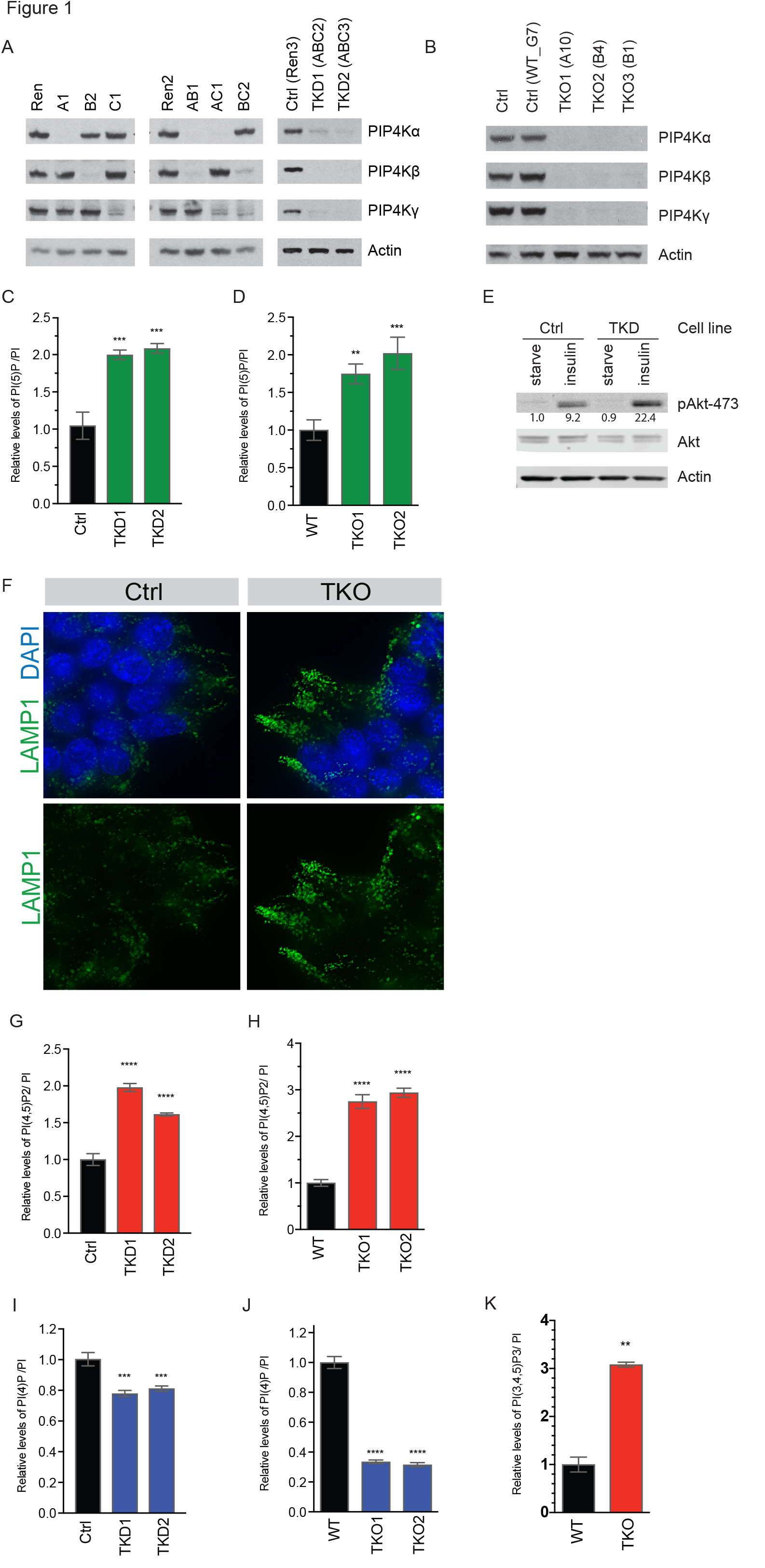
Validation of tools to eliminate all three PIP4K isoforms reveals paradoxical increase in PI(4,5)P_2_. (A) Western blots showing efficiency of knockdown of PIP4K isoforms in Hela cells. (B) Western blots showing 293T clones with CRISPR mediated knockout of *PIP4K2A, PIP4K2B*, and *PIP4K2C* (TKO cells). (C and D) Quantification by HPLC of PI(5)P which increases in cells with loss of PIP4K, shown in Hela TKD cells (C) as well as 293T TKO cells (D). (E) Knockdown of PIP4K enhances insulin signaling, as shown by western blot. Quantification by Licor indicated below respective bands. (F) Immunofluorescence detection of lysosomal marker Lamp1 in 293T control and TKO cells, representative images shown. (G and H) PI(4,5)P_2_ increases in cells with loss of PIP4K, shown in Hela TKD cells (G) as well as 293T TKO cells (H), measured by HPLC. (I and J) PI(4)P decreases in cells with loss of PIP4K, shown in Hela TKD cells (I) as well as 293T TKO cells (J), measured by HPLC. (K) 293T TKO cells have higher levels of PI(3,4,5)P_3_ as measured by HPLC. Significance calculated using ANOVA with Holm-Sidak multiple comparisons to control cell line with intact PIP4K. **p<0.01, ***p<0.001, ****p<0.0001

### Loss of PIP4Ks results in increased PI(5)P levels, increased PI3K activation, and defects in autophagy

PIP4K2A/B/C triple knockout 293T cells (hereafter TKO) and triple knockdown HeLa cells (hereafter TKD) recapitulated previously reported phenotypes observed when one or more of these genes were deleted or knocked down in other cell lines or in animals (Lamia et al., 2004; Lundquist et al., 2018; Emerling et al., 2013; Gupta et al., 2013). As expected, loss of all three isoforms conferred a two-fold increase in the substrate of these enzymes, PI(5)P (Figures 1C-D). In addition, TKD cells exhibited increased PI3K signaling upon insulin stimulation (Figure 1E), consistent with observations of increased insulin sensitivity in *Pip4k2b^-/-^* mice (Lamia et al., 2004). Furthermore, 293T TKO cells accumulated LAMP1 lysosomes, which we attributed to defects in autophagosome-lysosome fusion as previously described in *Pip4k2a^-/-,^ Pip4k2b^-/-^* mouse embryonic fibroblasts (Figure1F-G) (Lundquist et al., 2018).

### Cellular PI(4,5)P_2_ is increased upon depletion of PIP4K

Interestingly, depletion of the three PIP4Ks in cells altered the levels of phosphoinositides beyond PI(5)P. In both TKD and TKO cells, we observed a surprising increase in PI(4,5)P_2_ and concomitant decrease in PI(4)P (Figures 1H-K, Figures S2F-N). The increase in PI(4,5)P_2_ relative to PI(4)P is consistent with a more robust production of PI(4,5)P_2_ by the PIP5Ks. Additionally, although in the absence of acute stimulation, basal levels of PI(3,4,5)P_3_ are typically below the detection limit in HPLC-based assays, we were able to detect basal elevation of PI(3,4,5)P_3_ in TKO cells, suggesting that the increased PI(4,5)P_2_ is available to be utilized by cells in downstream signaling pathways (Figure 1L).

### Identification of a catalytic-independent function of PIP4K

Members of the PIP4K family have dramatically different catalytic rates, with PIP4Kα displaying 1000-fold higher activity than the near-enzymatically-dead PIP4Kγ (Clarke and Irvine, 2013). Analysis of PI(4,5)P_2_ levels in cells with single or double knockdown of PIP4K isoforms shows that there is an additive effect amongst all three isoforms that does not correlate with their relative catalytic activities (Figures 2A-B). Double knockdown of the most active isoforms, PIP4Kα/β, did not completely phenocopy the triple knockdown, suggesting that catalytic activity may not be the most important factor in suppressing PI(4,5)P_2_ levels. To evaluate whether the catalytic activity of the PIP4K enzymes is important for suppression of PI(4,5)P_2_ production, we generated shRNA-resistant PIP4K lentiviral vectors to express HA-tagged kinase-active or -dead PIP4K isoforms (Figures 2C-D).

**Figure 2.**
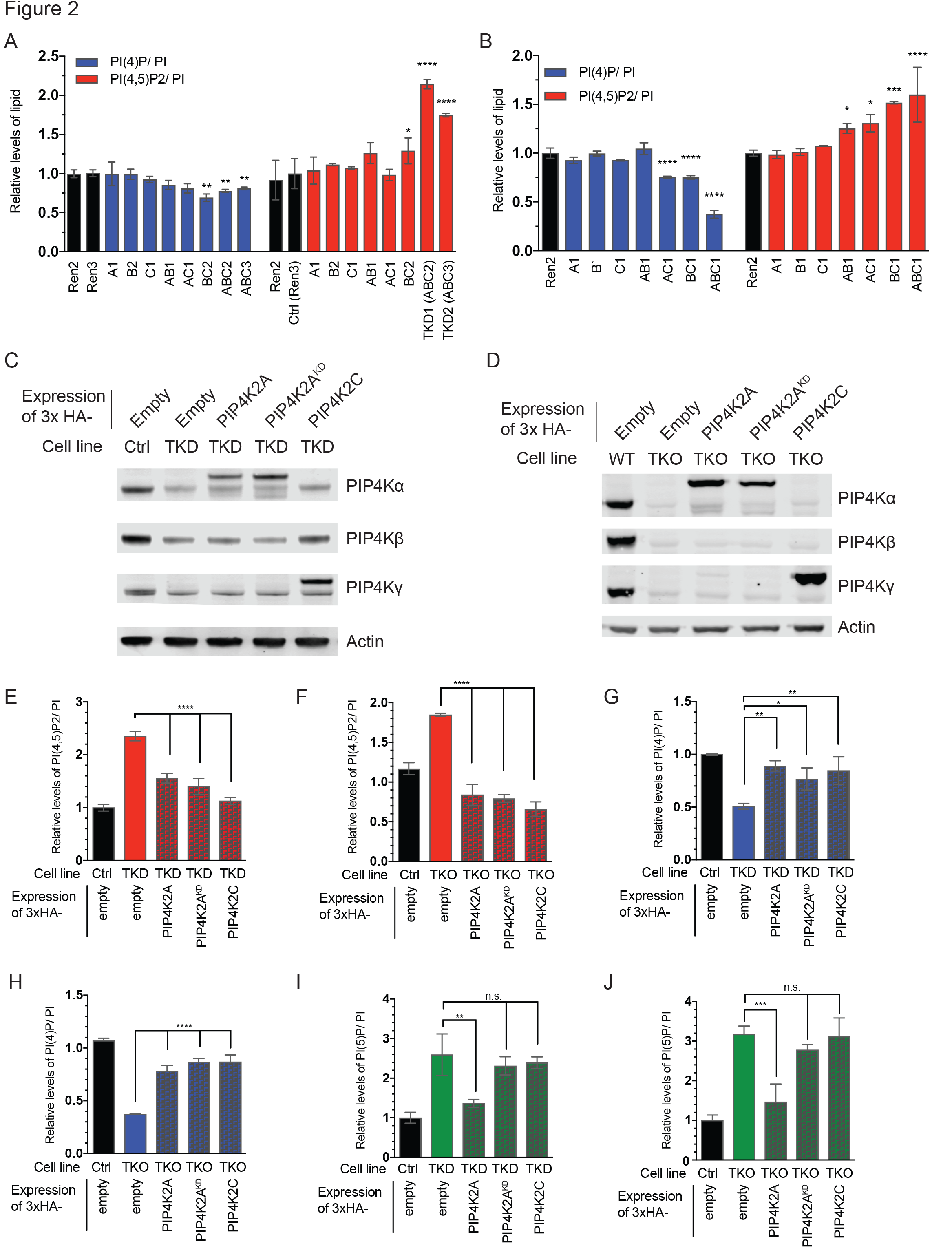
Identification of a catalytic-independent function of PIP4K. (A and B) Quantification of PI(4)P and PI(4,5)P2 levels by HPLC upon knockdown of one or multiple isoforms of PIP4K in 293T cells (A) or Hela cells (B) using tandem shRNA constructs. Hairpin constructs are indicated on y-axis. (C and D) Western blot validation of rescue cell line panel with reconstitution of active or kinase dead PIP4K2A^KD^ or active PIP4Kγ into Hela TKD cells or 293T TKO cells at near endogenous levels. (E-J) Quantification by HPLC of PI(4,5)P_2_ in Hela TKD rescue cell lines (E) and 293T TKO rescue cell lines (F). Quantification of PI(4)P in Hela TKD rescue cell lines (G) and 293T TKO rescue cell lines (H). Quantification of PI(5)P in Hela TKD rescue cell lines (I) and 293T TKO rescue cell lines (J). Significance calculated using ANOVA with Holm-Sidak multiple comparisons to TKD or TKO cell lines. *p<0.05, **p<0.01, ***p<0.001, ****p<0.0001

Surprisingly, expression of either kinase-active or -dead PIP4Ks reduced PI(4,5)P_2_ back to levels seen in parental cells (Figures 2E-F). Moreover, PI(4)P levels also returned to baseline with expression of either active or kinase-dead PIP4Kα (PIP4Kα^KD^) in both TKD and TKO cell lines (Figures 2G-H). In contrast, only the active PIP4Kα restored normal PI(5)P levels, indicating that the elevation of this lipid species in knockdown cells is due to the loss of PIP4K enzymatic activity (Figures 2I-J). From these observations, we conclude that PIP4Ks have a catalytic-independent role in maintaining homeostasis of cellular PI(4)P and PI(4,5)P_2_ levels in addition to their enzymatic function in converting PI(5)P to PI(4,5)P_2_.

### PIP5K activity is elevated in cells depleted of PIP4K

We performed a kinase assay using mechanically disrupted cells and found that TKO cells exhibit elevated PIP5K activity, consistent with the model that PIP5K is more active in the absence of PIP4K. The increased PIP5K activity was reversed in cells expressing either active or kinase-dead PIP4K isoforms, indicating it is not dependent on the enzymatic activity of these proteins (Figure 3A).

All three isoforms of PIP5K are stimulated by phosphatidic acid (PA) and association with G-proteins, such as Rac (Jenkins et al., 1994; Weernink et al., 2004). However, PIP5K activation in PIP4K depleted cells is not due to these upstream effectors as cells with PIP4K knockdown did not exhibit increased PA levels, and knockout of Rac1 did not abrogate the ability of PIP4K to modulate levels of PI(4)P and PI(4,5)P_2_ (Figure S3A-D). Furthermore, protein levels of PIP5K1A and PIP5K1C were not consistently altered in response to knockdown or knockout of the PIP4K family (Figure S3E-F).

**Figure 3.**
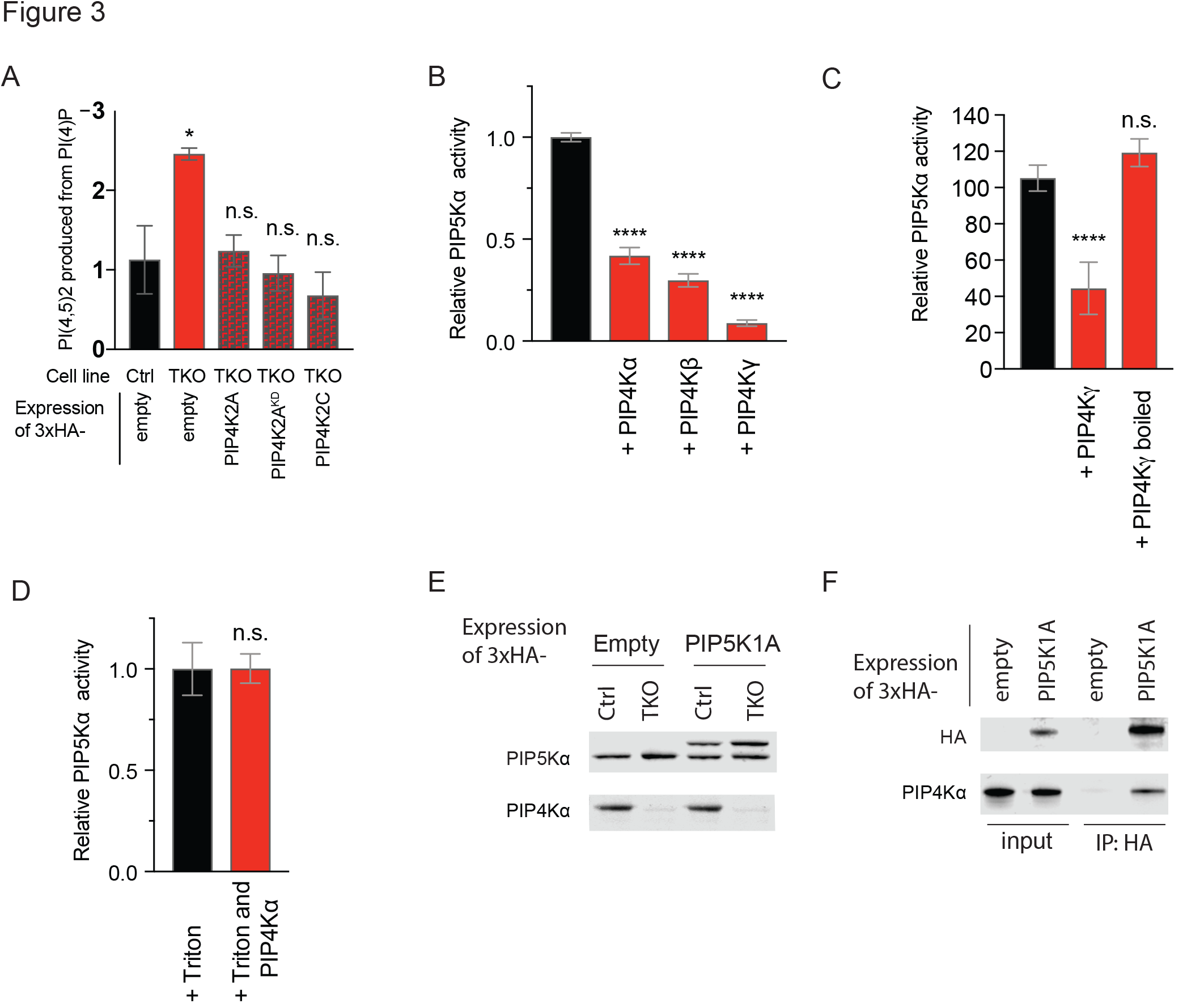
PIP4K inhibits PIP5K activity. (A) Changes in PIP5K activity in 293T cells. Cell pellets were normalized to cell number and sonicated in the presence of excess PI(4)P. Radioactive ATP was added for kinase reaction. Lipids were extracted and deacylated for quantification of PI(4,5)P_2_ levels by HPLC. (B) PIP5Kα activity is inhibited *in-vitro* by addition of PIP4Kα, PIP4Kβ, or PIP4Kγ. Significance calculated using ANOVA with Holm-Sidak multiple comparisons to control cell line. *p<0.05, **p<0.01, ***p<0.001, ****p<0.0001, n.s. not significant (C) Quantification of *in vitro* kinase assay measuring conversion of PI(4)P to radiolabeled PI(4,5)P_2_ by PIP5Kα. PIP4Kγ was added in native or denatured state. Significance calculated using ANOVA with Holm-Sidak multiple comparisons to control cell line. ****p<0.0001, n.s. not significant (D) Kinase reaction the presence of 0.1% Triton-X100, PIP4Kα can no longer inhibit PIP5Kα activity *in vitro*. (E) Western blot validation of near-endogenous level expression of 3xHA-tagged PIP5Kα levels in 293T ctrl and TKO cells. (F) Co-immunoprecipitation of PIP4Kα with HA-PIP5Kα from 293T TKO cells expressing empty 3xHA vector or 3xHA-tagged PIP5Kα.

### PIP4K can inhibit PIP5K through direct interactions on the surface of negatively charged membranes

Members of the PIP4K and PIP5K family have been previously reported to form complexes with each other (Hinchliffe et al., 2002). Additionally, unbiased, high-throughput mass spectrometry has revealed evidence of interaction between PIP5Kα and PIP4Kα (Huttlin et al., 2017). Such an interaction would be an opportunity for PIP4K to directly inhibit PIP5K. Notably, PIP4K family members are considerably more abundant than PIP5K family members, making the stoichiometric ratio favorable for PIP4Ks to inhibit a substantial pool of PIP5Ks (Itzhak et al., 2016; Wisniewski et al., 2014). To test whether the altered PIP5K activity in cells depleted of PIP4Ks could be a result of a disrupted physical interaction between these two families of kinases, we performed *in vitro* kinase reactions with purified proteins. Using 5- to 10-fold molar excess of PIP4Ks, we found that all three PIP4K isoforms could directly inhibit the PIP5Kα-catalyzed phosphorylation of PI(4)P into PI(4,5)P_2_ (Figure 3B). To eliminate the possibility that a small molecule contaminant from our *E. coli* PIP4K purifications could inhibit PIP5Kα, we boiled PIP4Kγ and found that denatured PIP4Kγ could no longer inhibit PIP5Kα (Figure 3C). To investigate if the interactions between PIP4K and PIP5K occur at the membrane bilayer, we repeated the experiment with detergent to eliminate the formation of PI(4)P-containing liposomes. Presenting phosphoinositides in Triton-X100 micelles enhances the activity of some PI kinases, such as the PI4Ks (Guo et al., 2003), but we found that it reduces PIP5Kα activity overall. Importantly, addition of detergent eliminates the ability of PIP4Kα to inhibit PIP5Kα (Figure 3D). These data suggest that recruitment of both proteins to the membrane surface is required to mediate the inhibitory effect. As we performed this assay using vast molar excess of the substrate PI(4)P relative to the levels of both enzymes, we postulate that sequestration of PI(4)P by PIP4Ks is unlikely to account for this inhibition. Attempts to co-immunoprecipitate PIP4K with PIP5K in the presence of detergent were unsuccessful, which is not surprising given the membrane-dependence of the association of PIP4K and PIP5K. However, by chemical crosslinking of proteins in intact cells, we demonstrated interaction of 3xHA-tagged PIP5Kα expressed at near-endogenous levels with endogenous PIP4Kα (Figures 3E and 3F), supporting the hypothesis that PIP4Ks interact with PIP5Ks in cells to regulate PI(4,5)P_2_ levels.

These results reveal a novel and potentially general mechanism for regulation of PI(4,5)P_2_ production, wherein PIP5K is negatively regulated by local presence of PIP4Ks. As discussed above, protein copy numbers of PIP4K family members are in 10 to 50-fold excess to that of PIP5K family members across a panel of cell lines, well in excess of the amount needed to inhibit PIP5K *in vitro* (Wisniewski et al., 2014). We propose that a fraction of cellular PIP4Ks are directly interacting with PIP5K enzymes at membranes enriched in PI(4)P, and functions to attenuate the PIP5K-mediated conversion of PI(4)P to PI(4,5)P_2_. Though, historically, attempts to directly visualize the subcellular localization of PIP4Ks have been inconsistent, recent global quantitative mapping of protein subcellular localization suggest that both PIP5Ks and PIP4Ks are largely bound to plasma membranes (Itzhak et al., 2016).

### Structural role of PIP4K in regulating PIP5K and PI3K pathway is distinct from its catalytic role in autophagy

Levels of PI(4,5)P_2_ are remarkably stable to perturbations in the metabolism of its precursor PI(4)P (Nakatsu et al., 2012), yet we find that depletion of the PIP4K family members results in nearly two-fold changes in PI(4)P and PI(4,5)P_2_, as previously shown. Insulin activation of the PI3K pathway is initiated at the plasma membrane, where conversion of PI(4,5)P_2_ into PI(3,4,5)P_3_ recruits effector proteins Akt and Pdk1 to propagate signaling cascades. We hypothesized that, in PIP4K-depleted cells, activation of the PI3K pathway could be enhanced due to increased availability of its substrate, PI(4,5)P2. HPLC analysis of lipids showed that PIP4K knockout cells exhibited a five-fold increase in PI(3,4,5)P_3_ upon acute insulin stimulation (Figure 4A). Consistent with the model that PIP4Ks suppress insulin signaling by their catalytic-independent suppression of PI(4,5)P_2_ synthesis by the PIP5Ks, expression of the kinase-dead PIP4K2A or the inherently low-activity enzyme PIP4Kγ in TKO cells was able to reverse the insulin-dependent increase in PI(3,4,5)P_3_ accumulation.

**Figure 4.**
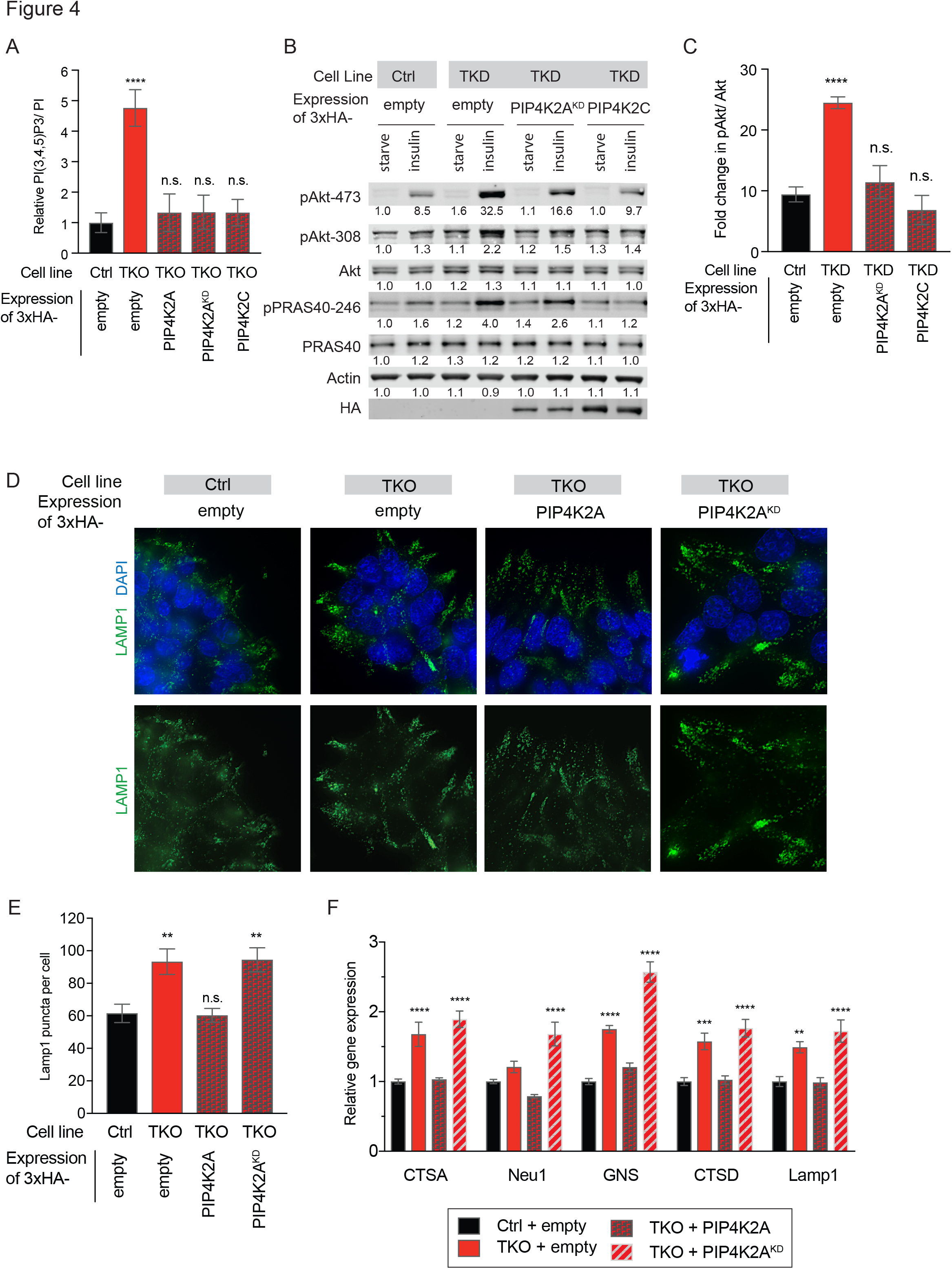
Structural role of PIP4K in regulating PIP5K and the PI3K pathway is distinct from its catalytic role in autophagy. (A) Insulin stimulation of PI(3,4,5)P_3_ synthesis by PI3K in 293T rescue cell lines, quantified by HPLC. Cells were serum starved for 20 hours, then stimulated with 50 ng/mL insulin for 5 minutes (n=4). (B) Insulin stimulation of Akt activation in Hela rescue cell line panel. Cells were serum starved for 12 hours, then stimulated with 250 ng/mL insulin for 10 minutes. Western blot band intensities were measured by Licor and are indicated under the relevant band. (C) Insulin-stimulated increase in pAkt-473, normalized to total protein, across multiple experiments. (D) Immunofluorescence detection of lysosomal marker Lamp1 in 293T rescue cell line panel. Representative images are shown. (E) Quantification of Lamp1 puncta per cell in 293T rescue cell line panel. (F) qRT-PCR for TFEB target genes in rescue cell line panel. Significance calculated using ANOVA with Holm-Sidak multiple comparisons to control cell line. *p<0.05, **p<0.01, ***p<0.001, ****p<0.0001, n.s. not significant

Further, HeLa TKD cells displayed enhanced PI3K pathway activation, as judged by S473 phosphorylation of AKT as well as AKT-mediated phosphorylation of PRAS40 (Manning and Toker, 2017) (Figures 4B-C). Expression of either PIP4Kα^KD^ or PIP4Kγ reversed the enhanced insulin sensitivity whether measured by PI(3,4,5)P_3_ levels or AKT activation. Our results indicate that increasing PIP5K activity by eliminating PIP4K isoforms drives overproduction of PI(4,5)P_2_ in plasma membrane pools that are used by PI3K during insulin signaling.

Finally, we asked whether the novel non-catalytic function of PIP4Ks was required for the previously reported role of PIP4K2A in mediating autophagy (Lundquist et al., 2018). As expected, loss of PIP4Ks resulted in accumulation of LAMP1-positive lysosomes and increased TFEB transcriptional targets. However, unlike the changes observed in lipid species, only reconstitution with active, but not kinase-dead, PIP4K2A showed restored regulation of autophagy (Figures 4D-F). Together, this work highlights the complexity and multi-functionality of PIP4K enzymes which has not previously been appreciated.

## DISCUSSION

In this study, we show that the proteins encoded by the *PIP4K2A, PIP4K2B* and *PIP4K2C* genes have distinct catalytic and non-catalytic functions and provide insight into phenotypes observed in mice in which these genes have been deleted. The roles of the PIP4K2A and PIP4K2B isoforms in autophagosome-lysosome fusion that was recently reported in mouse embryonic fibroblasts and *in vivo* using mice with targeted liver-specific PIP4K deletions (Lundquist et al., 2018), clearly depends on the ability of these enzymes to convert PI(5)P to PI(4,5)P_2_. However, our current study revealed that the increased insulin sensitivity previously observed in *Pip4k2b^-/-^* mice (Lamia et al., 2004) is likely due to loss of a non-catalytic function of PIP4K2B in suppressing the ability of PIP5Ks to produce PI(4,5)P_2_, the substrate for PI 3-kinase (Figure S3I).

We show that PIP4Ks and PIP5Ks directly interact and while this is a likely mechanism for inhibition, other indirect mechanisms may also explain the observed suppression of PIP5K activity. Because PIP4Ks are highly abundant, in excess of PIP5Ks, and bound to negatively charged membranes, these enzymes could alter local membrane structures or recruit other enzymes to modify PIP5Ks to modulate their activities. Future studies will address these possibilities.

Our results indicate that design of drugs to promote degradation of PIP4Ks may mitigate insulin resistance, and possibly retard progression of type II diabetes. Recently emerging technologies raise the possibility that one could degrade specific PIP4K proteins by linking them to E3 ligases (Bondeson and Crews, 2017). We initially had reservations about development of such drugs due to embryonic lethality of germline co-deletion of PIP4K2B/2C or PIP4K2A/2B (Emerling et al., 2013; Shim et al., 2016).

However, our results suggest that most cell lines do not require PIP4K isoforms for viability, at least when grown in tissue culture with full nutrients. Embryonic lethality from co-deletion of murine PIP4K isoforms may be attributed to early developmental defects unique to mammalian development, and partial and/or tissue-specific targeted depletion may be well tolerated.

We expect that each tissue has a different ratio of PIP4Kα: PIP4Kβ: PIP4Kγ, and therefore, it remains to be seen if targeting a subset of the PIP4K family will enhance systemic insulin sensitivity. Whereas the PIP4Kβ knockout mice exhibit increased insulin sensitivity, this is not seen in PIP4Kα and PIP4Kγ knockout mice, and may be because PIP4Kβ is the predominant isoform in skeletal muscle, which is a primary site of insulin-mediated glucose uptake. It remains to be seen if other insulin-responsive tissues, such as liver and adipose tissue, will become sensitized to insulin when multiple PIP4K isoforms are depleted.

PIP4K family members repress the conversion of PI(4)P to PI(4,5)P_2_ (and subsequently to PI(3,4,5)P_3_) to limit PI3K pathway signaling. This occurs by a mechanism that does not require the catalytic activity of these enzymes, while they directly catalyze the conversion of PI(5)P to PI(4,5)P_2_ to promote autophagy. How did these two disparate functions come to reside in the same protein? One possibility is to ensure that levels of PI(4)P remain high at local sites of autophagosome-lysosome fusion. Several recent studies indicate that PI(4)P is required for autophagosome-lysosome fusion and that, in some cases, this PI(4)P is derived from an inositol 5-phosphatase acting on local pools of PI(4,5)P_2_ (De Leo et al., 2016; Wang et al., 2015). Thus, the PIP4Ks may be both producing the local PI(4,5)P_2_ at the autophagosome-lysosome junction and ensuring that, upon conversion to PI(4)P, it is not converted back to PI(4,5)P_2_ by a PIP5K. This model would ensure a unidirectional conversion of PI(5)P to PI(4)P and efficient autophagosome-lysosome fusion. The partial localization of PIP4Kγ to the Golgi (Clarke et al., 2008) could also suppress conversion of Golgi PI(4)P to PI(4,5)P_2_ to maintain high local concentration of the PI(4)P needed for vesicle trafficking at this location.

In the future, it will be important to characterize how accumulation of PI(4,5)P_2_ in PIP4K-depleted cells affects other cellular processes, such as cell adehesion, migration and/or receptor tyrosine kinase signaling. Further, it will be essential to examine how this change impacts the localization and levels of other phosphoinositides within cells.

In all, our findings highlight an unexpected and important separation of catalytic and non-catalytic functions in PIP4K family enzymes and indicate new avenues for intervention in enhancing insulin signaling.

## ACKNOWLEDGEMENTS

We thank all members of the Cantley Laboratory for feedback. We thank Jared Johnson for chemistry and biochemistry consultations. We thank Yuxiang Zheng for sharing expertise on HPLC analysis of phosphoinositides. We thank Hyeseok Shim for insight on PIP4K2C biology. This work was supported by grants to from the NIH (R35 CA197588, U54 U54CA210184) and the Lustgarten Foundation, and grants to J.M.B., including NIH Pathway to Independence (R00 GM110121), NSF CAREER (CHE-1749919), and Beckman Young Investigator (Arnold and Mabel Beckman Foundation). to L.C.C. LED was supported by a K22 Career Development Award from the NCI/NIH (CA 181280-01). D.G.W. was supported by a Medical Scientist Training Program grant from the NIH (T32GM007739) to the Weill Cornell/Rockefeller/Sloan-Kettering Tri-Institutional MD-PhD Program, and T.W.B. was supported by an NSF graduate research fellowship (DGE-1650441).

## AUTHOR CONTRIBUTIONS

D.G.W., M.N.P, L.E.D., and L.C.C. formulated the research plan and interpreted experimental results. D.G.W, M.N.P, M.L, designed and performed the experiments with assistance from J.S., OM., S.A, A.H, M.F. L.E.D., B.H., T.W.B., J.M.B., and T.J.Y. provided reagents and helped interpret experimental results. D.G.W, M.N.P, and L.C.C. wrote the manuscript and all authors edited it.

---

**SUPPLEMENTAL INFORMATION**

**Figure S1.**
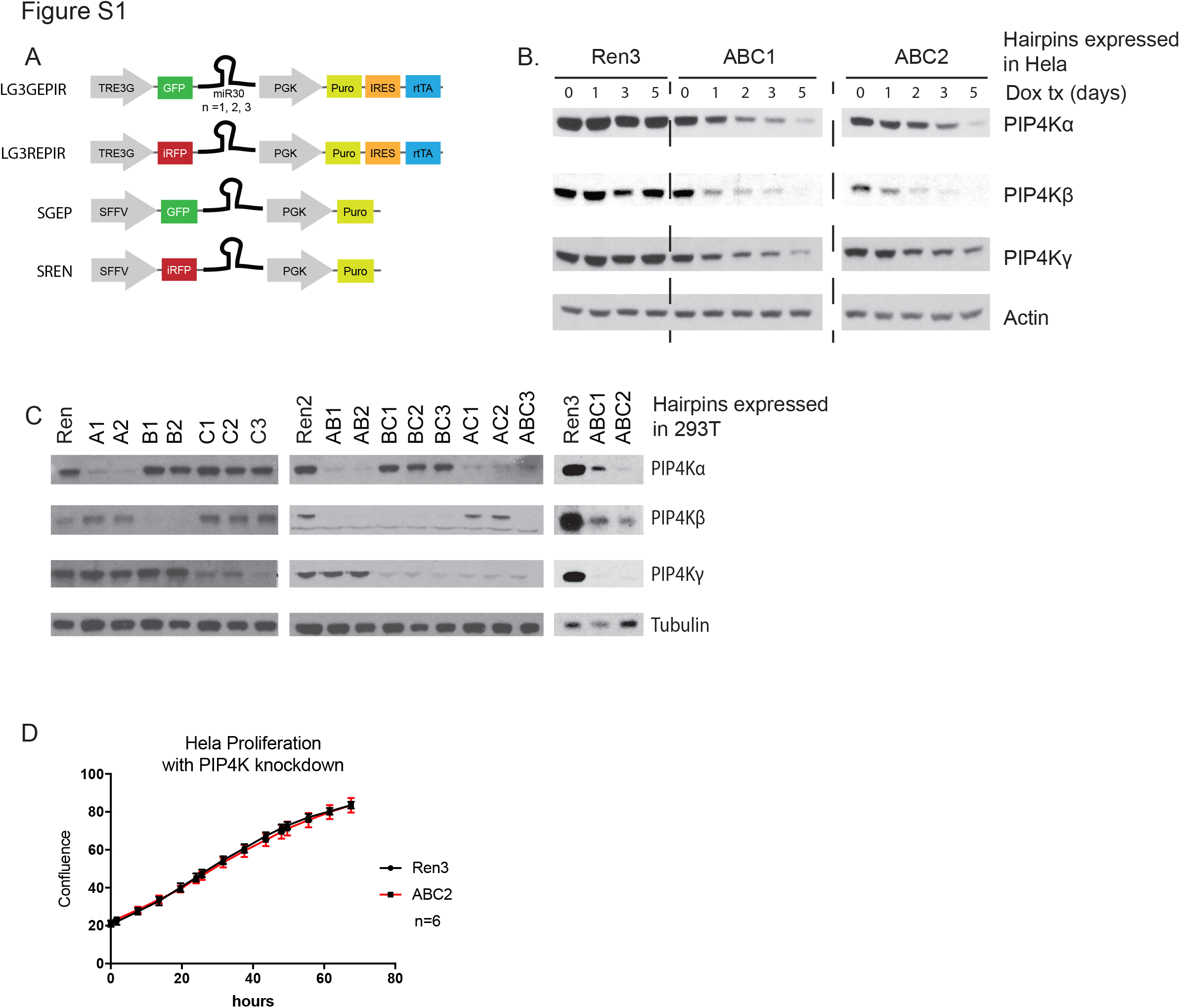
Knockdown of Multiple PIP4K Isoforms, Related to Figure 1. (A) Schematic of miRE expression cassette in doxycycline inducible vectors (LG3GEPIR, LG3REPIR) or constitutive (SGEP, SREN). Hairpin expression can be concatemerized with GFP or infrared RFP (iRFP). (B) Kinetics of shRNA mediated knockdown of all three isoforms of PIP4K. 293Ts expressing doxycycline-inducible hairpins were harvested over the course of five days of doxycycline treatment. (C) Validation of knockdown of PIP4K isoforms in 293T cells. (D) Cell proliferation over 72 hours, graphed as mean confluence with SEM. N=6

**Figure S2.**
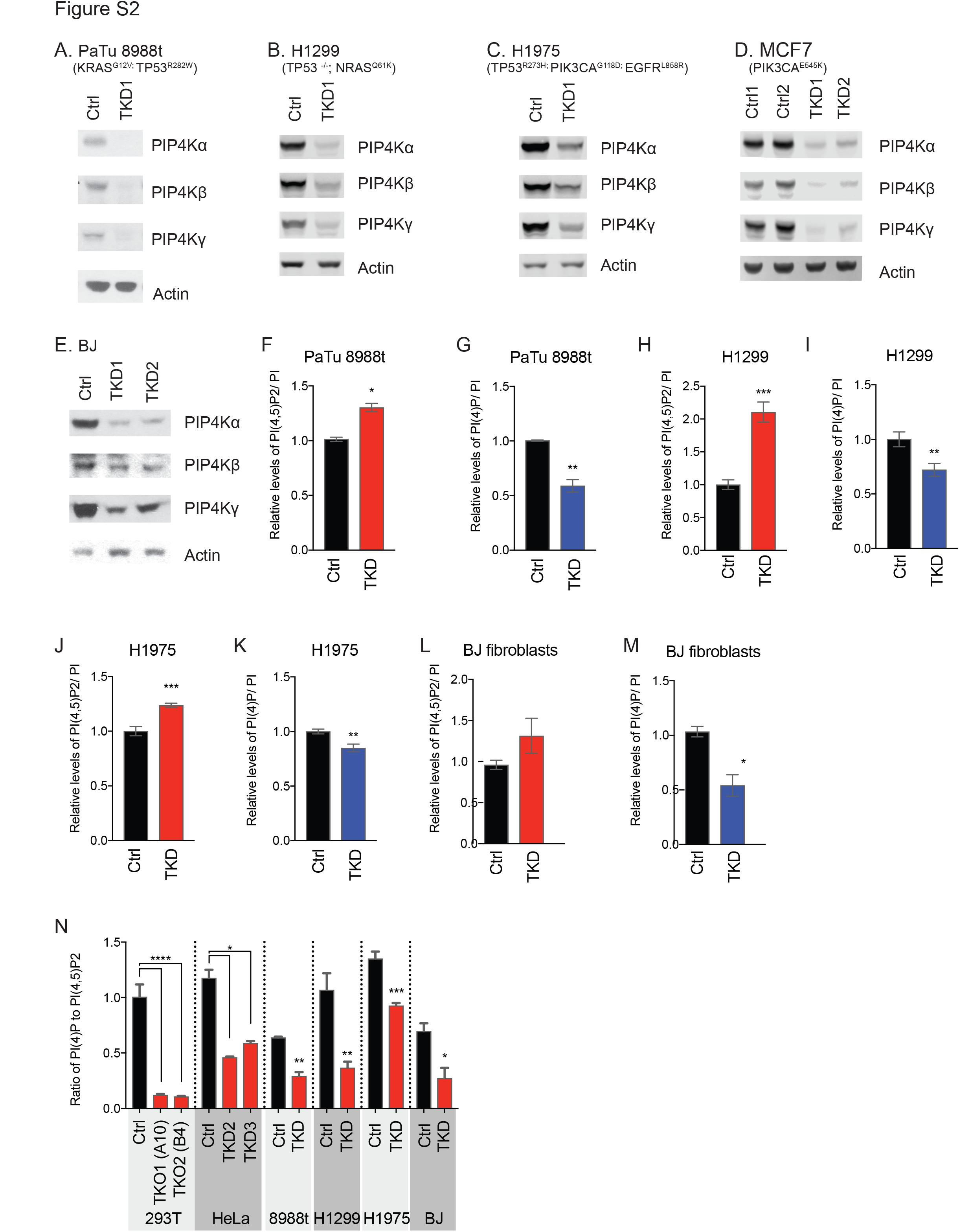
Depletion of PIP4K in a Panel of Cell Lines of Carious Tissue and Mutational Backgrounds, Related to Figure 1. (A-E) Knockdown of PIP4Ks in a panel of cell lines. Mutations reported from Cancer Cell Line Encyclopedia (CCLE). (F-M) Quantification of PI(4,5)P_2_ and PI(4)P levels in cell lines, n=3. (N) Quantification of PI(3,4,5)P_3_ in 293T control versus TKO cells during exponential growth in full media. Significance calculated using ANOVA with Holm-Sidak multiple comparisons to control cell line. *p<0.05, **p<0.01, ***p<0.001

**Figure S3.**
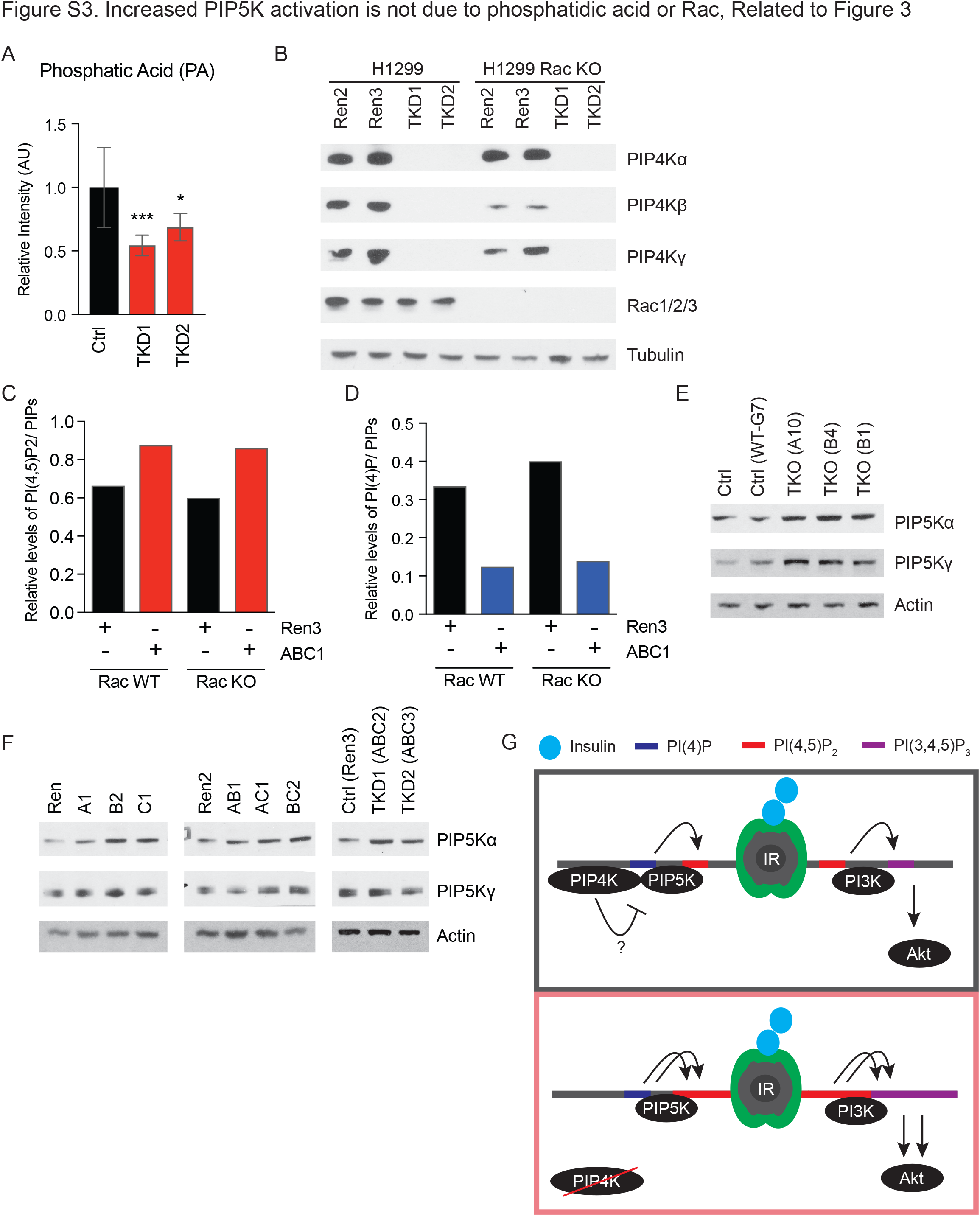
Increased PIP5K Activation is Not Due to Phosphatidic Acid or Rac, Related to Figure 3. (A) Phosphatidic acid quantified in Hela TKD cells. Samples were normalized to content of phosphatidylethanolamine. Analysis performed in triplicate. Significance calculated using ANOVA with Holm-Sidak multiple comparisons to control cell line. *p<0.05, ***p<0.001 (B) Validation of H1299 Rac KO cells with triple knockdown of PIP4K isoforms. (C and D) Measurement of PI(4,5)P_2_ (C) and PI(4)P (D) levels in H1299 Rac KO cells. Lipids were normalized to total phosphorylated lipids rather than PI content. N=1 (E and F) Levels of PIP5Kα and PIP5Kγ in 293T TKO clones (E) as well as Hela knockdown cell lines (F). (G) At the plasma membrane, loss of PIP4K de-represses PIP5K, thereby causing increased PI(4,5)P_2_ production and increased PI(3,4,5)P_3_ production during insulin stimulation.

## METHODS

### CONTACT FOR REAGENT AND RESOURCE SHARING

Further information and requests for resources and reagents should be directed to and will be fulfilled by the Lead Contact, Lewis C. Cantley.

### EXPERIMENTAL MODEL AND SUBJECT DETAILS

#### Cell lines, authentication

Cell lines were purchased from ATCC and/or fingerprinted with the University of Arizona genetics core.

#### Cell culture conditions

293T, HeLa, PaTu 8988t, and BJ cells were cultured using DMEM media supplemented with 10% FBS, glutamine and pyruvate. H1299 and H1975 cells were cultured in RPMI media.

### METHOD DETAILS

#### Cell lysis and immunoblotting

Cells were lysed in RIPA buffer supplemented with 1 tablet of protease and phosphatase inhibitor. After incubation on ice for 20 minutes, lysates were cleared by centrifugation at 14,000 x g and supernatant was quantified using BCA assay. Lysates were subjected to SDS-PAGE using Novex NuPAGE system. Proteins were separated on 4-12% Bis Tris Pre-Cast Gels 10% Bis Tris gels using MOPS buffer. Proteins were transferred to 0.45 μm nitrocellulose membranes at 350 mA for 1h. Membranes were blocked in 5% non-fat milk in TBST and incubated with primary antibody overnight. For chemi-luminescently detect antibody binding, membranes were blotted with HRP conjugated secondary antibodies. Membranes were developed using ECL solution, and exposed to film. For insulin signaling westerns, protocol was modified such that cells were lysed in triton buffer, IRDye secondary was used for LiCor Odyssey detection with quantification using Image Studio Lite software.

#### Generation of lentivirus, viral transduction

293T cells were used to generate lentivirus. Once cells were at 90-95% confluence in a 10cm dish, transfection was performed with opti-mem, lipofectamine 2000, lentiviral vector, and accessory plasmids VSVG and ∆8.2. Virus supernatant was harvested 2 days and 3 days post transfection, then filtered and concentrated in an ultracentrifuge at 25,000 rpm for 120 minutes at 4°C.

#### Generation of cell lines with CRISPR knockout of PIP4K

CRISPR guides in pX458 were transfected in 293T cells. At 48-96 hours post transfection, GFP positive cells were single-cell sorted in 96-well plates using the Influx sorter at the WCMC Flow Cytometry Core. Two weeks later, wells were scored to contain single cell colony and expanded to screen for successful PIP4K2A/ PIP4K2B/ PIP4K2C knockout. Validation was performed by western blotting as well as PCR around each cut site.

#### Generation of cell lines with miRE knockdown of PIP4K

LT3GEPIR vectors containing desired miR-E shRNA(s) were double digested downstream of existing shRNA(s) with EcoRI-HF (NEB R31010) and MluI-HF (NEB 3198) and PCR purified. The miR-E sequence to be added was PCR amplified with Multi-sh-fw (5’-agg cgcgaagactcaattgaaggctcgagaaggtatattgctg-3’) and Multi-sh-rev (5’-cacttttttcaattgacacgtacgcgtattctaccgggta-3’). PCR products were purified, double digested with BbsI/MluI (NEB R0539, R0198, buffer 2.1), and PCR purified once more. Ligations were performed with PCR product and open LT3GEPIR vectors using T4 ligase. Colonies were screened using miRE-fwd.

#### Generation of cell lines with stable expression of cDNAs

To generate vectors to express hairpin-resistant coding sequences for PIP4K isoforms, we cloned wild type PIP4K2A and PIP4K2C into a lentiviral backbone with a PGK promoter. Next, we generated mutations in PIP4K cDNA using QuikChange (Agilent 200522) to make kinase-dead variants and silent mutations in wobble positions for hairpin-resistant cDNA. No wobble mutations were needed for PIP4K2C cDNA since all hairpins targeted the 3’ UTR.

#### Measurement of phosphoinositides with high performance liquid chromatography

Cellular phosphoinositides were metabolically labeled for 48 hours in inositol-free DMEM supplemented with glutamine,10% dialyzed FBS, and 10 μCi/ mL 3H myo-inositol. Cells were washed with PBS and then transferred on ice. Cells were killed and then harvested by scraping using 1.5 mL ice-cold aqueous solution (1M HCl, 5 mM Tetrabutylammonium bisulfate, 25 mM EDTA). 2 mL of ice cold MeOH and 4mL of CHCl3 were added to each sample. After ensuring each vial is tightly capped, samples were vortexed and then centrifuged at 1000 rpm for 5 min. If a significant intermediate layer was visible, sample were gently agitated and spun again, until there were predominately two clear layers. The organic layer (lower) was cleaned using theoretical upper, while the aqueous layer was cleaned using theoretical lower (theoretical upper and lower made by combining CHCl3: MeOH: aqueous solution in 8:4:3 v/v ratio). Organic phases were collected and dried under nitrogen gas. Lipids were deacylated using monomethylamine solution (47% Methanol, 36% of 40% Methylamine, 9% butanol, and 8% H2O, by volume). Samples were incubated at 55° for 1 hour and subsequently dried under nitrogen gas. To the dried vials, 1 mL of theoretical upper and 1.5 mL of theoretical lower were added (theoretical upper and lower made by combining CHCl3:MeOH:H2O in 8:4:3 v/v ratio). Samples were vortexed and spun at 1000rpm. The aqueous phase (upper) was collected and dried under nitrogen gas. Samples were resuspended in 150 μL Buffer A (1mM EDTA), filtered and transferred to Agilent polypropylene tubes. Samples were analyzed by anion-exchange HPLC using Partisphere SAX column. The compounds were eluted with a gradient starting at 100% Buffer A (1mM EDTA) and increasing Buffer B (1mM EDTA, 1M NaH2PO4) over time: 0-1 min 100% Buffer A, 1-30 min 98% Buffer A/2% Buffer B, 30-31 min 86% Buffer A/14% Buffer B, 31-60 min 70% Buffer A/ 30% Buffer B, 60-80 min 34% Buffer A/ 66 % Buffer B, 80-85 min 100% Buffer B, 85-120 min 100% Buffer A. Buffers were pumped at 1 mL/min through column. Eluate from the HPLC column flowed into an on-line continuous flow scintillation detector for isotope detection. The detector was set to observe events between 10 minutes and 85 minutes, with scintillation fluid flowing at 4 mL/min.

#### Measurement of phosphatidic acid using LCMS

Cellular lipids were extracted using same method described for phosphoinositide analysis. Lipids were dried and diluted with 113 μL of chloroform:methanol:water (73:23:3) mixture and filtered (0.45 μm) before analysis on an Agilent 6230 electrospray ionization–time-of-flight (ESI–TOF) MS coupled to an Agilent 1260 HPLC equipped with a Phenomenex Luna silica 3 μm 100 Å 5 cm x 2.0 mm column. LCMS analysis was performed using normal phase HPLC with a binary gradient elution system where solvent A was chloroform:methanol:ammonium hydroxide (85:15:0.5) and solvent B was chloroform:methanol:water:ammonium hydroxide (60:34:5:0.5). Separation was achieved using a linear gradient from 100% A to 100% B over 9 min. Phospholipid species were detected using a dual ESI source operating in positive mode, acquiring in extended dynamic range from m/z 100–1700 at one spectrum per second; gas temperature: 325 °C; drying gas 10 L/min; nebulizer: 20 psig; fragmentor 300 V

##### Immunoprecipitation

293T cell lines expressing 3xHA empty vector or 3xHA-PIP5K1A were fixed for 10 minutes in 4% PFA, washed 3x with PBS, and lysed in lysis buffer (10mM Tris 7.4, 150 mM NaCl, 0.5 mM EDTA, 0.5% NP40) using probe sonicator for 2 minutes. Lysates were pre-cleared with control magnetic beads for 1 hour and 4° and Pierce magnetic anti-HA beads added for overnight incubation at 4°. Beads were washed 3x with lysis buffer and aliquots were added to sample buffer loading dye with beta mercaptoethanol, boiled, and run on SDS-page for immunoblotting.

#### Fluorescence microscopy

293T cells were grown on glass coverslips pre-treated with poly-d-lysine. Adherent cell lines were rinsed with phosphate-buffered saline, pH 7.4 (PBS) and subsequently fixed/permeabilized in -20 MeOH for 20 minutes. After fixation, the cells were blocked for 30 minutes in blocking buffer (PBS with 3% BSA) and labeled with primary antibodies in blocking buffer for 1 hour at room temperature. Coverslips were washed three times with blocking buffer and incubated with Alexa Fluor-conjugated goat secondary antibodies in blocking buffer for 1 hour at room temperature. After incubation with secondary antibodies, coverslips were washed three times with PBS, once with water, and then mounted on a glass microscope slide with Prolong Gold with DAPI. The following primary antibodies were used: LAMP1. Alexa Fluor-conjugated secondary antibodies were used at 1:1000. Fluorescent and phase contrast images were acquired on a Nikon Eclipse Ti microscope equipped with an Andor Zyla sCMOS camera. Within each experiment, exposure times were kept constant and in the linear range throughout. When using the 60x oil immersion objectives, stacks of images were taken and deconvoluted using AutoQuant.

#### qRT-PCR

Total RNA was prepared using RNeasy. cDNA was synthesized using Superscript Vilo and qRT-PCR performed utilizing Fast SYBR green and the Realplex Mastercycler. For a list of primers used see oligos in resource table. Isolation of mRNA and qPCR was performed as follows. 200,000 cells were plated in 6-well plastic dishes. 24 hours later, the RNA in the lysates was extracted using the RNeasy protocol. The RNA was resuspended in 50 μl H_2_O at a concentration of 1 μg/μL. cDNA was transcribed using the SuperScript Vilo. The sequences of the oligonucleotides used as primers in the PCR reactions are given in Table S4. The genes that were quantified here were previously shown to be regulated by TFEB

##### In-vitro kinase assays

Cells were trypsinized and normalized for cell number. Pellets were resuspended in HNE buffer (20 mM HEPES pH 7.4, 100mM NaCl, 0.5mM EGTA), and sonicated in the presence of 32P-γ-ATP, and liposomes (4ug PS, 2 ug PI(4)P in 30mM HEPES pH 7.4, 1mM EGTA). Reactions were stopped by added 50uL of 4N HCL. To extract lipids, 100 μL of MeOH:CHCl3 (1:1) was added and samples were vortexed 2x 30 seconds. Samples were spun down at top speed for 2 min and the organic phase containing phosphatidylinositol lipids (bottom) were separated using thin-layer chromatography (TLC) using 1-propanol: 2N acetic acid (65:35 v/v). TLC plates were prepared ahead of time by coating with 1% Potassium Oxalate. Phosphorylated lipids were visualized by autoradiography on a GE Typhoon FLA 7000 and quantified using ImageQuant TL software.

### QUANTIFIICATION AND STATISTICAL ANALYSIS

Experiments were repeated with at least three biological replicates with the following exceptions: experiments in Figure S3 were performed once, and panels with error bars were performed with technical triplicates. No samples were excluded from analysis. When comparing two groups, a two-tailed t-test was used. P values are indicated in figure legends. When comparing greater than two groups, significance was calculated using ANOVA and Holm-Sidack post-hoc test. Figure legends indicate which comparisons are significant, with respective p-values.

**Table S1.**
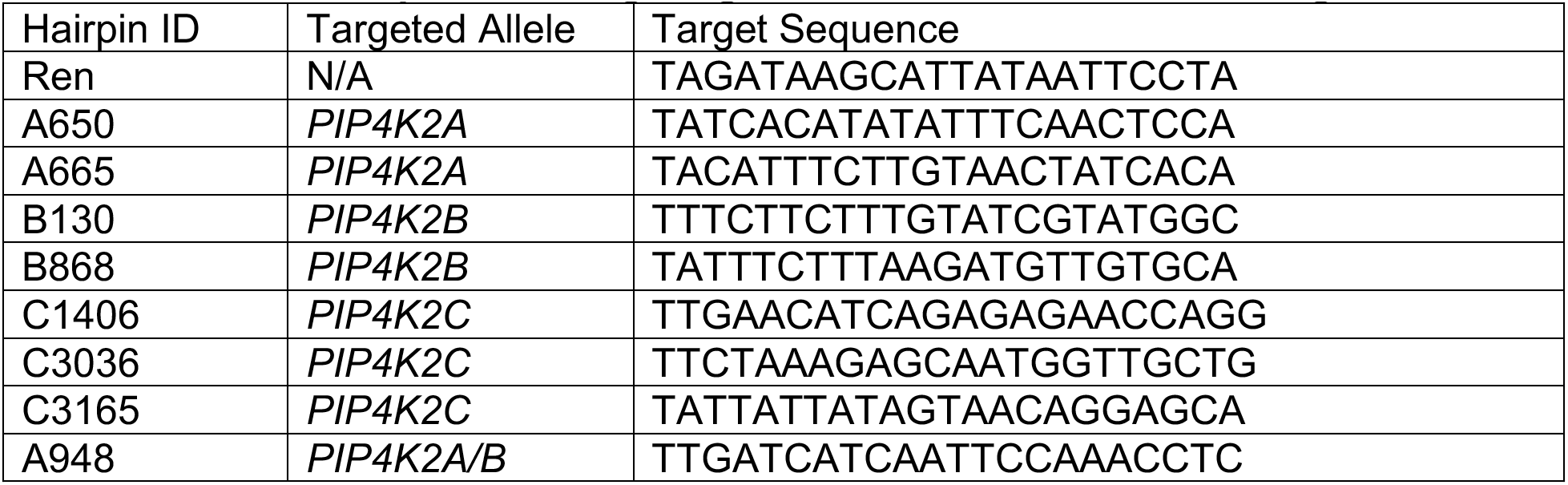
shRNA Sequences Targeting PIP4K Isoforms, Related to Figure 1.

**Table S2.** Optimized Tandem shRNA series, Related to Figure 1.

| Hairpin Code | Tandem Hairpin IDs |
| --- | --- |
| Ren | Ren |
| Ren2 | Ren; Ren |
| Ren3 | Ren; Ren; Ren |
| A1 | A650 |
| A2 | A665 |
| B1 | B130 |
| B2 | B868 |
| C1 | C1406 |
| C2 | C3036 |
| AB1 | A650; B868 |
| AB2 | A665; B130 |
| BC1 | B130; C3165 |
| BC2 | B868; C1406 |
| AC1 | A650; B868 |
| AC2 | A665; B130 |
| TKD1 (ABC1) | A650; B130; C1406 |
| TKD2 (ABC2) | A665; B868; C3036 |
| TKD3 (ABC3) | A948; C3165 |

**Table S3.**
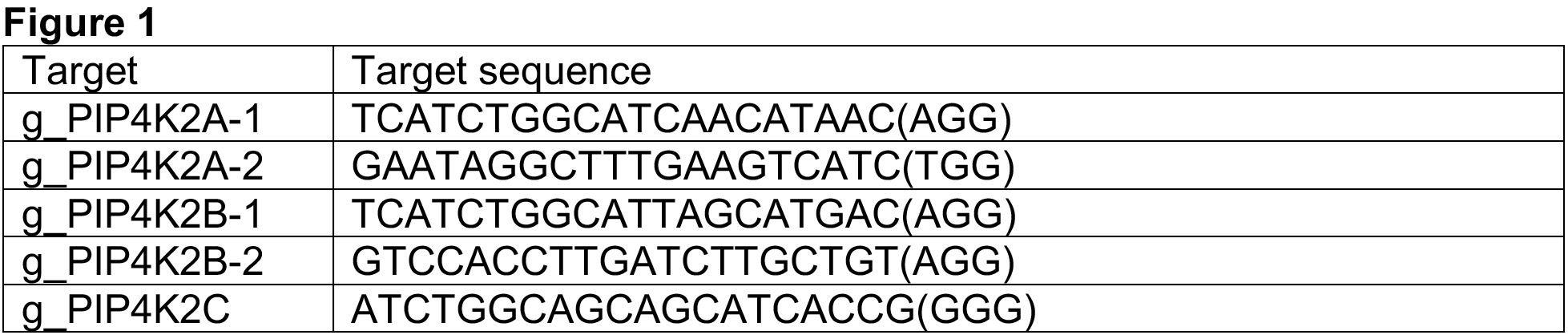
CRISPR Guides Targeting *PIP4K2A, PIP4K2B, PIP4K2C*, Related to Figure 1.

